# Boosting detection of low abundance proteins in thermal proteome profiling experiments by addition of an isobaric trigger channel to TMT multiplexes

**DOI:** 10.1101/2020.12.30.424894

**Authors:** Sarah A. Peck Justice, Neil A. McCracken, José F. Victorino, Aruna B. Wijeratne, Amber L. Mosley

**Affiliations:** Department of Biochemistry and Molecular Biology, Indiana University School of Medicine, Indianapolis, Indiana 46202, United States

## Abstract

The study of low abundance proteins is a challenge to discovery-based proteomics. Mass-spectrometry (MS) applications, such as thermal proteome profiling (TPP) face specific challenges in detection of the whole proteome as a consequence of the use of nondenaturing extraction buffers. TPP is a powerful method for the study of protein thermal stability, but quantitative accuracy is highly dependent on consistent detection. Therefore, TPP can be limited in its amenability to study low abundance proteins that tend to have stochastic or poor detection by MS. To address this challenge, we incorporated an affinity purified protein complex sample at submolar concentrations as an isobaric trigger channel into a mutant TPP (mTPP) workflow to provide reproducible detection and quantitation of the low abundance subunits of the Cleavage and Polyadenylation Factor (CPF) complex. The inclusion of an isobaric protein complex trigger channel increased detection an average of 40x for previously detected subunits and facilitated detection of CPF subunits that were previously below the limit of detection. Importantly, these gains in CPF detection did not cause large changes in melt temperature (T_m_) calculations for other unrelated proteins in the samples, with a high positive correlation between T_m_ estimates in samples with and without isobaric trigger channel addition. Overall, the incorporation of affinity purified protein complex as an isobaric trigger channel within a TMT multiplex for mTPP experiments is an effective and reproducible way to gather thermal profiling data on proteins that are not readily detected using the original TPP or mTPP protocols.

---

Proteins are the functional units of a cell, carrying out and controlling processes at specific times and locations to maintain homeostasis and respond to external stimuli. As a consequence of functional changes, proteins can exist in a variety of biophysical states within cells as a consequence of variants in their primary sequence, post-translational modification (PTM) state, and/or subcellular localization. In many cases a protein’s biophysical state is impacted by associations with other proteins, including both transient and stable protein-protein interactions. The characterization of protein-protein interactions (PPIs) is fundamental to gaining a full understanding of biological mechanism. In fact, PPIs are so critical to proper protein function that disruptions in these interactions often lead to disease and/or cell death ^1^. Advances in mass spectrometry (MS)-based proteomics workflows continue to increase our ability to study protein complex dynamics and PPIs^2–7^. MS-based approaches for protein interaction analysis rely on discovery-based proteomics performed using data-dependent acquisition (DDA). Generally in DDA, peptides with the most intense ions from MS1 are selected for fragmentation and MS2 analysis ^8^. This approach maximizes signal to noise levels and thereby increases confidence in the selection and subsequent identification of the peptide ions.

Challenges with the use of DDA include selection of peptide ions from protein(s) of interest that are present at low relative abundance levels or when peptides of interest (such as PTM containing peptides) are present at low relative levels to their unmodified counterparts. Low abundance peptides may be present at insufficient MS1 signal intensity levels to trigger fragmentation and MS2 analysis based on instrument settings for MS2 analysis. While fractionation and an extended HPLC gradient help to spread out the elution of peptides into the mass spectrometer, many peptides may still co-elute such that highly abundant ion species will outcompete those that are less abundant ^9^. A number of strategies have recently emerged to improve MS detection of low abundance proteins and post-translational modifications (PTMs) for a variety of applications including single cell proteomics^10–16^. Although we will not discuss all of the recently established strategies here, one such strategy, Boosting to Amplify Signal with Isobaric Labeling (BASIL), has similarities that have informed the current work. Specifically, BASIL has been shown to successfully increase detection of low abundance phosphopeptides through addition of a boosting sample to a tandem mass tag (TMT)-based multiplex^17^. TMTPro labeling allows for the multiplexing and relative quantitation of up to 16 samples^18–20^. As each TMT label is isobaric, labeled peptides from the multiplexed samples elute into the mass spectrometer together and are analyzed simultaneously as one ion peak during MS1 scans which is distinguished in fragment ion scans during MSn (typically MS2 or MS3) analysis. By incorporating a phospho-enriched sample into a single channel in the TMT multiplex, Yi *et al* increased ion abundance of phosphopeptides in the MS1 scan to the extent that MS2 was triggered for phosphopeptides that were typically below the level of detection in standard DDA approaches ^17^. BASIL allowed for the identification and quantification of phosphopeptides in other TMT channels where enrichment had not been performed^17^. The BASIL method has since been optimized for detection of phosphopeptides in single cells^21^ and similar approaches have been applied to phosphotyrosine-containing peptides^22^, SILAC-labeled peptides^23^, and using synthetic peptides to particular peptides of interest^24^. BASIL and other similar methods that take advantage of isobaric carrier channels could have numerous applications in DDA-based quantitative workflows.

The challenges to studying low abundance proteins in DDA proteomics experiments extend in particular to the mass spectrometry-based thermal proteome profiling (TPP) methods and are the focus of this study. TPP analysis takes advantage of TMT labeling technology to produce protein melt curves that can then be compared across conditions to measure alterations in protein thermal stability^25, 26^. Although TPP was originally developed to study drug and ligand binding, it has been shown to also be a robust approach to probe PPIs in a number of different applications (recently reviewed by Mateus *et al* ^27^). We recently developed a new application of TPP referred to as mutant TPP (mTPP), that is used to study the effects of protein missense mutations on the proteome at large with the ability to focus in on specific protein complexes and their PPIs^28^. mTPP analysis is advantageous to other methods for the study of PPIs in that it does not require antibodies, addition of reagents such as crosslinkers, or the genetic manipulations (such as the production of fusion proteins) typically necessary for many other PPI analyses. Additionally, mTPP can be performed with significantly less starting material than traditional affinity purification or enrichment approaches, making it applicable to a wider variety of sample types. Despite these advantages, we have quickly encountered challenges associated with quantitative analysis of specific target proteins and their interaction partners. Therefore, a strategy for increasing the ion intensity of proteins of interest in mTPP experiments would have a significant impact on our ability to study PPI dynamics of low abundance protein complexes while still retaining the context of changes within the overall proteome. One advantage of TMT- and iTRAQ-based multiplexed workflows for global proteomics studies is that the pooling of multiple samples generates increased protein starting material that can then be subjected to extensive biochemical fractionation to facilitate deep proteome coverage ^29–33^. This advantage can be coupled with protein extraction methods using denaturants such as urea or SDS to isolate the full proteome of many cells and tissues ^34^. The workflow for TPP cannot exploit these advantages since: 1) Temperature treatment of lysates for TPP results in unequal levels of protein mixture across the multiplex that, in our hands, vary on average at least 10-fold from the lowest to the highest temperature treatment ^28^; and 2) Non-denaturing protein extraction buffers must be used to maintain protein structure, PPIs, and protein interactions with other molecules (including but not limited to lipids, metabolites, small molecules, and drugs) ^25–27^. As a consequence, TPP workflows typically result in decreased proteome coverage relative to denaturant extracted proteomes even when equivalent amounts of starting material are used ^28^.

To expand proteome coverage for our mTPP workflow, we have developed a BASIL-like approach to increase the signal of low abundance protein complexes and their representative peptides in mTPP experiments using a protein complex affinity purification trigger channel in place of the phosphopeptides isobaric boosting channel used in BASIL^17^. As a proof-of-concept, we investigated the ability of this approach to enhance detection of the relatively low abundance protein complex cleavage and polyadenylation factor (CPF) complex in a mTPP workflow. Affinity purified CPF that we have previously characterized ^35–39^ was incorporated as an isobaric trigger channel into our mTPP workflow at a ratio to the lowest heat-treated mTPP sample of ~1:8 and ~1:50. Using this approach, a significant increase in the abundance of CPF complex members was observed, including those that were not readily identified without the isobaric trigger channel. Importantly, addition of an isobaric trigger channel into our mTPP workflow does not appear to have a significant impact on the melt temperature (T_m_) calculation of proteins detected both with and without the trigger. Overall, the use of an isobaric trigger channel is a robust approach to prioritize DDA selection of proteins or peptides of interest such as missense mutant containing proteins and their interaction partners, which are of particular focus within mTPP experiments.

## EXPERIMENTAL SECTION

### Yeast strains and growth

All experiments were performed in *Saccharomyces cerevisiae.* The parental strain SMY732, described previously,^40^ was obtained from the Mirkin lab and used in the trigger experiments comparing technical replicates. For the biological replicate experiments, the wildtype strain used was the commercially available *BY4741* strain (Open Biosystems). The *ssu72-2* temperature sensitive mutant (first described by the Hampsey lab ^41^) was purchased from Euroscarf. The Pta1-FLAG strain was made via homologous recombination. The 3×FLAG tag DNA sequence was amplified from plasmids obtained from Funakoshi and Hochstrasser ^42^ to insert the FLAG epitope tag into the genome at the 3’-end of the *PTA1* gene in WT (BY4741). Successful incorporation of the FLAG tag was confirmed via Western blot.

For mTPP experiments, cells were inoculated at an OD_600_ = 0.3 and grown to an OD_600_ = 0.8 in yeast extract, peptone, dextrose (YPD) medium at permissive temperature (30°C or 25°C). YPD was removed by filtration through a nitrocellulose membrane (Millipore, Burlington, MA). Cells were flash frozen with liquid nitrogen and stored at −80°C to be used in subsequent sample preparation steps. For affinity purification of CPF via Pta1-FLAG, cells were grown overnight at 30°C in YPD to an OD_600_ ~3. Cells were pelleted, washed, and transferred to 50ml conical tubes for storage at −80° until subsequent sample preparation steps.

### Sample preparation

BY4741 and *ssu72-2* samples for mTPP were prepared as described in Peck Justice et al^28^ with the exception of an extended temperature range for the heat treatment. For the no trigger mTPP experiments, lysate was treated at the following ten temperatures: untreated, 25°, 35°, 46.2°, 48.8°, 51.2°, 53.2°, 55.2°, 56.5°, and 74.9°C. A TMT 10plex kit (Thermo Scientific, Waltham, MA) with channels TMT126; TMT127N; TMT127C; TMT128N; TMT128C; TMT129N; TMT129C; TMT130N; TMT130C and TMT131 were respectively used to label peptide solutions derived from untreated, 25°, 35°, 46.2°, 48.8°, 51.2°, 53.2°, 55.2°, 56.5°, and 74.9°C temperature treatments in WT. In *ssu72-2*, channels TMT126; TMT127N; TMT127C; TMT128N; TMT128C; TMT129N; TMT129C; TMT130N; TMT130C and TMT131 were respectively used to label peptide solutions derived from untreated, 25°, 35°, 48.8°, 46.2°, 51.2°, 74.9°C, 53.2°, 55.2°, and 56.5° temperature treatments. TMT labeling steps were performed according to manufacturer provided instructions.

To boost detection of the native CPF subunits, subsequent mTPP replicates of WT and *ssu72-2* included the addition of a trigger channel consisting of an affinity-purified CPF complexes. Affinity purification of CPF via Pta1-FLAG was performed as described previously for Ssu72-FLAG purifications ^35^. The Pta1-FLAG affinity purified sample was added at a ratio of 6.25 ug trigger to 50 ug of the lowest heat-treated sample (1:8 ratio) for the initial study. The untreated samples were removed from the multiplex from no trigger samples to accommodate for the isobaric trigger channel to be labeled with TMT126. The remainder of the channels, TMT127N; TMT127C; TMT128N; TMT128C; TMT129N; TMT129C; TMT130N; TMT130C and TMT131 were used to label peptide solutions derived from 25°, 35°, 46.2°, 48.8°, 51.2°, 53.2°, 55.2°, 56.5°, and 74.9°C temperature treatments. Subsequent sample preparation steps were performed as described in Peck Justice *et al*^28^.

SMY732 samples for independent replicate experiments were prepared as described in Peck Justice *et al*^28^. Lysate was treated at the following eight temperatures: 25°, 35°, 48.8°, 51.2°, 53.2°, 55.2°, 56.5°, and 74.9°C. A TMT 16plex kit (Thermo Scientific, Waltham, MA) with channels TMT127N; TMT127C; TMT128N; TMT128C; TMT129N; TMT129C; TMT130N; TMT130C were respectively used to label peptide solutions derived from 25°, 35°, 48.8°, 51.2°, 53.2°, 55.2°, 56.5°, and 74.9°C temperature treatments in parental culture samples. Note that some channels in the 16plex were used for other samples not described in this report. These heat-treated lysates were analyzed twice and as separate LC-MS experiments for comparison of technical replicate reproducibility. In one experiment, the set of combined labeled samples was analyzed with a ninth trigger channel (TMT126) at a ratio of 1 ug total isobaric trigger channel protein to 50 ug of the lowest heat-treated sample (1:50 ratio) which included the Pta1-FLAG affinity purified material (described previously) while in the second experiment, the trigger was not added.

### LC-MS/MS analysis

Following multiplex preparation as described above, samples were subjected to high-pH reversed phase fractionation as previously described ^28^. NanoLC-MS/MS analyses were performed on an Orbitrap Fusion Lumos mass spectrometer (Thermo Scientific, Waltham, MA) coupled to an EASY-nLC HPLC (Thermo Scientific, Waltham, MA). One-third of the resuspended fractions were loaded onto an in-house prepared reversed phase column using 600 bar as applied maximum pressure to an Easy-Nano 25cm column with 2μm reversed phase resin. The peptides were eluted using a 180-minute gradient increasing from 95% buffer A (0.1% formic acid in water) and 5% buffer B (0.1% formic acid in acetonitrile) to 25% buffer B at a flow rate of 400 nL/min. The peptides were eluted using a 180-minute gradient increasing from 95% buffer A (0.1% formic acid in water) and 5% buffer B (0.1% formic acid in acetonitrile) to 25% buffer B at a flow rate of 400 nL/min. Nano-LC mobile phase was introduced into the mass spectrometer using a Nanospray Source (Thermo Scientific, Waltham, MA). During peptide elution, the heated capillary temperature was kept at 275°C and ion spray voltage was kept at 2.6 kV. The mass spectrometer method was operated in positive ion mode for 180 minutes having a cycle time of 4 seconds for MS/MS acquisition. MS data was acquired using a data-dependent acquisition using a top speed method following the first survey MS scan. During MS1, using a wide quadrupole isolation, survey scans were obtained with an Orbitrap resolution of 120 k with vendor defined parameters—m/z scan range, 375-1500; maximum injection time, 50; AGC target, 4E5; micro scans, 1; RF Lens (%), 30; “DataType”, profile; Polarity, Positive with no source fragmentation and to include charge states 2 to 7 for fragmentation. Dynamic exclusion for fragmentation was kept at 60 seconds. During MS2, the following vendor defined parameters were assigned to isolate and fragment the selected precursor ions. Isolation mode = Quadrupole; Isolation Offset = Off; Isolation Window = 0.7; Multi-notch Isolation = False; Scan Range Mode = Auto Normal; FirstMass = 120; Activation Type = CID; Collision Energy (%) = 35; Activation Time = 10 ms; Activation Q = 0.25; Multistage Activation = False; Detector Type = IonTrap; Ion Trap Scan Rate = Turbo; Maximum Injection Time = 50 ms; AGC Target = 1E4; Microscans = 1; DataType = Centroid. During MS3, daughter ions selected from neutral losses (e.g. H_2_O or NH_3_) of precursor ion CID during MS2 were subjected to further fragmentation using higher-energy C-trap dissociation (HCD) to obtain TMT reporter ions and peptide specific fragment ions using following vendor defined parameters. Isolation Mode = Quadrupole; Isolation Window =2; Multi-notch Isolation = True; MS2 Isolation Window (m/z) = 2; Number of notches = 3; Collision Energy (%) = 65; Orbitrap Resolution = 50k; Scan Range (m/z) = 100-500; Maximum Injection Time = 105 ms; AGC Target = 1E5; DataType = Centroid. The data were recorded using Thermo Scientific Xcalibur (4.1.31.9) software (Copyright 2017 Thermo Fisher Scientific Inc.).

### Protein Identification and Quantification

Resulting RAW files were analyzed using Proteome Discoverer™ 2.4 (Thermo Scientific, Waltham, MA). The SEQUEST HT search engine was used to search against a yeast protein database from the UniProt sequence database (December 2015) containing 6,279 yeast protein and common contaminant sequences (FASTA file used available on ProteomeXchange under accession PXD020689). Specific search parameters used were: trypsin as the proteolytic enzyme, peptides with a max of two missed cleavages, precursor mass tolerance of 10 ppm, and a fragment mass tolerance of 0.02 Da. Static modifications used for the search were, 1) carbamidomethylation on cysteine residues; 2) TMTsixplex label on lysine (K) residues and the N-termini of peptides. Dynamic modifications used for the search were oxidation of methio-nine and acetylation of N-termini. Percolator False Discovery Rate was set to a strict setting of 0.01. Values from both unique and razor peptides were used for quantification. No normalization setting was used for protein quantification since the different temperature treatments are expected to have different protein amounts. The mass spectrometry proteomics data have been deposited to the ProteomeXchange Consortium via the PRIDE^43^ partner repository with the dataset identifier PXD020689 and doi: 10.6019/PXD020689.

### Data analysis

Venn Diagrams were created using Venny 2.1^44^. Dot plots, scatter plots, and waterfall plots were created using ggplot2^45^ in R Studio (R Studio for Mac, Version 1.2.5001). Bar graphs were created in Excel (Microsoft Excel for Mac, Version 16.38). The TPP package (v3.12.0)^46^ in R Studio was used to generate normalized melt curves and to determine protein melt temperatures as described previously^26^. Resulting data processing and analysis also occurred in R Studio. Change in T_m_ (ΔT_m_) values were calculated by taking WT T_m_ -*ssu72-2* T_m_, thereby limiting calculations to proteins detected in both WT and mutant. Further parsing was accomplished by limiting our data to melt curves with r^2^ values > 0.9 and then by proteins that were detected in at least two of the three replicates. Proteins were ranked according to median change in T_m_ and ordered from the largest change (proteins that were destabilized in the mutant) to smallest change (proteins that were stabilized in the mutant). Changes in T_m_ that were outside of ± 2***σ*** (***σ*** being the standard deviation), were considered statistically significant, and identified as proteins destabilized or stabilized due to the mutations in *SSU72*.

## RESULTS AND DISCUSSION

### Addition of an affinity purified isobaric trigger channel to mTPP multiplexes does not cause large changes in peptide coverage or quantitation

We hypothesized that incorporation of a well-characterized affinity purified sample isolated from our system of interest as an isobaric trigger channel would increase MS1 ion intensity of peptides of interest within the TMT multiplex. As a consequence, the identification of peptides from the affinity purified protein complex would boost the identification in the remaining experimental mTPP channels used for melt curve production and subsequent T_m_ calculation when comparing different experimental samples. Similar to the approach used in BASIL^17^, the incorporation of an affinity purified CPF complex purified from our system of interest has numerous potential advantages including native levels of CPF post-translational modifications and protein interaction partners. Similar to mTPP, the affinity purifications for the CPF complex were performed using non-denaturing buffers to preserve PPIs. Qualitatively, the MS/MS fragment data for CPF complexes will be improved from inclusion of the isobaric trigger channel increasing the ion abundance of the fragments and therefore the probability of CPF identification at the peptide spectrum match (PSM) level. From a quantitative perspective, TMT126 information will be obtained during data processing but will be excluded for interpretation of the mTPP melt curves for each protein.

Pta1-3xFLAG affinity purifications were digested with LysC/Trypsin and labeled with TMT126 for inclusion within the mTPP multiplex. mTPP quantitative analysis and curve generation was performed using the remaining channels as described in the methods (Fig. 1). The mTPP samples were subjected to eight or nine different temperatures (25°, 35°, 46.2°, 48.8°, 51.2°, 53.2°, 55.2°, 56.5°, and 74.9°C) and then centrifuged to separate soluble and insoluble material as previously described ^28^. For samples with eight temperature points no 46.2° treatment sample was included. Samples were then processed and subjected to LC-MS/MS analysis using an MS2-based fragmentation and TMT quantitation workflow (Fig. 1). Using SEQUEST HT and Proteome Discoverer 2.4 for qualitative and quantitative analysis, between 1,750 and 3,150 proteins were detected and quantified depending on the replicate (Supp. Tab. 1). Replicates are designated as preparation 1, 2, 3 (hence p1, p2, p3). The p1 replicate had less IDs overall but p2 and p3 had very similar peptide detection levels (Supp. Tab. 1). To gain insights into general trends with the quantitative data, dot plots were generated to show the abundance value for each quantified protein (Fig. 2). Consistent with previous mTPP experiments^28^, there was an overall decrease in protein abundance as the temperature at which the sample was treated increased. Importantly, incorporation of a protein complex isobaric trigger channel into the multiplex did not alter the overall trend of decreasing protein abundance with increased temperature (Figure 2B&D) or have a significant effect on the number of proteins detected. The average ion abundance at each temperature treatment also remained consistent between samples plus or minus the isobaric trigger channel (compare Figure 2A to B and C to D). Finally, the average quantitative ratio of the isobaric trigger channel to the mTPP experimental sample processed at 25°C remains consistent at a 1:50 (Figure 2B) or 1:8 (Figure 2D) mirroring the ratios used for mixing of the multiplex.

**Figure 1.**
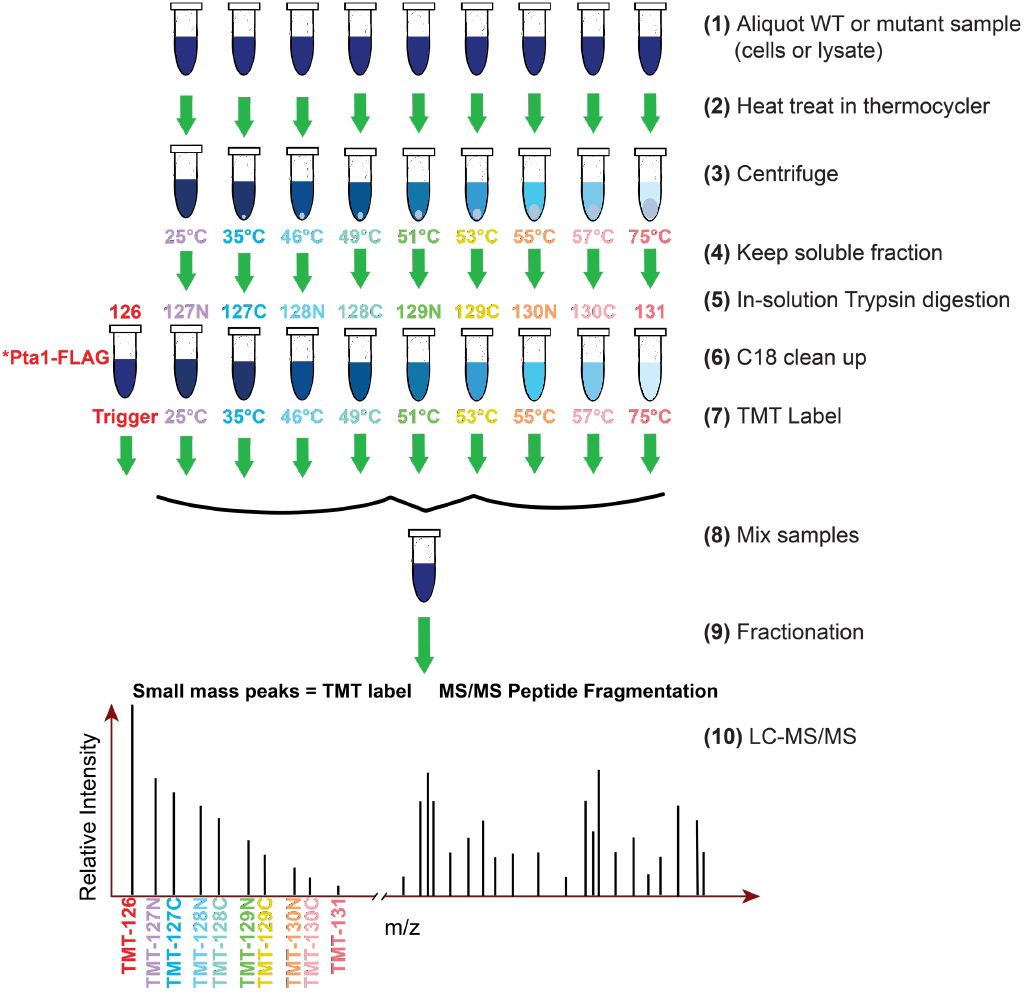
Workflow overview for mTPP with isobaric trigger channel addition. Equal amounts of protein from each lysate for every biological replicate sample were subjected to different temperature treatments: 25°, 35°, 46.2°, 48.8°, 51.2°, 53.2°, 55.2°, 56.5°, and 74.9°C, to induce protein denaturation. The soluble fractions from each treatment as well as a Pta1-FLAG affinity purification sample were digested in-solution with Trypsin/Lys-C. Resulting peptides were labeled with isobaric mass tags (TMT 10plex) as shown and mixed prior to mass spectrometry (MS) analysis. Resulting MS/MS data were analyzed using Proteome Discoverer™ 2.4 to identify and quantify abundance levels of peptides for each temperature treatment and each biological replicate across genotypes.

**Figure 2.**
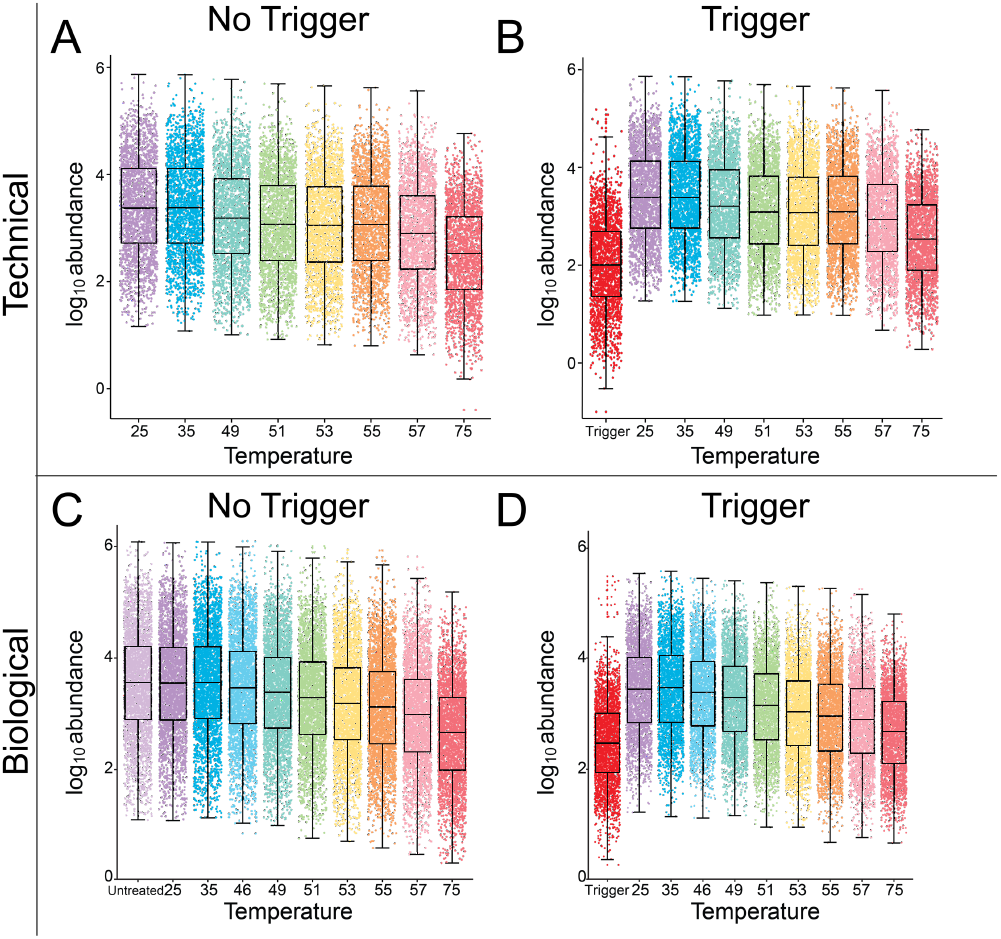
The use of an isobaric trigger channel does not alter mTPP experimental channel abundance values. Dot plots of protein abundance values for each protein detected in WT cells in technical replicates without (A) and with (B) the isobaric trigger channel (trigger) addition and representative biological replicates without (C) and with (D) the isobaric trigger addition. The same general decrease of protein abundances with increase in temperature treatment is seen across all replicates. Dot plots for additional replicates are provided in Supp. Fig. 1.

The impact of the trigger on mTPP analysis was investigated using both technical replicates and biological replicates so that we could evaluate differences in our workflow and their impact on qualitative and quantitative parameters. For the technical replicates, the same labeled samples were split into two TMT multiplexes; one multiplex without an isobaric CPF trigger (no trigger) and one multiplex with an isobaric CPF trigger labeled with TMT126 (trigger) with a quantitative ratio (based on protein assays) to lowest temperature treatment of ~1:50. For the biological replicates, four biological replicate samples were grown and prepared independently of one another. One replicate contained a non-heat treated (untreated) sample that was labeled with TMT126 (no trigger sample) and the remaining three replicates were multiplexes with a CPF trigger labeled with TMT126 (trigger) with a trigger to lowest temperature treatment ratio of ~1:8.

While there was not an obvious effect on the overall abundance of proteins in the samples, it is possible that the trigger could affect the detection and identification of proteins by biasing the mass spectrometer towards proteins present in the affinity purification. Comparisons of MS-based measurements across the technical replicates showed that the trigger channel incorporation did not have a significant impact on protein identification and quantification (Fig. 3A). Technical replicate analyses showed very similar numbers of detected PSMs, peptides, and proteins suggesting that the addition of the trigger channel at a ratio of 1:50 has little impact on overall LC-MS/MS detection (Fig. 3A, yellow). The biological replicates showed more variation across samples which is attributed to their separate processing for TPP in addition to variation that could occur from trypsin digestion and other processing steps ^47, 48^. Trigger p1 in the biological replicate study did have overall lower levels of proteins detected but this was not likely a consequence of trigger channel addition considering that Trigger p2 and Trigger p3 samples had similar detection levels to the No trigger sample (Fig. 3A, green). Direct comparison of proteins quantified in the No trigger vs. Trigger samples showed an 80% overlap in quantified proteins with unique proteins present in all individual datasets (Figure 3B&C). Overall, these data suggest that the addition of an isobaric trigger channel has little to no impact on overall proteome detection outside of the inherent variability seen in independent sample processing (for the biological replicates) and LC-MS/MS runs.

**Figure 3.**
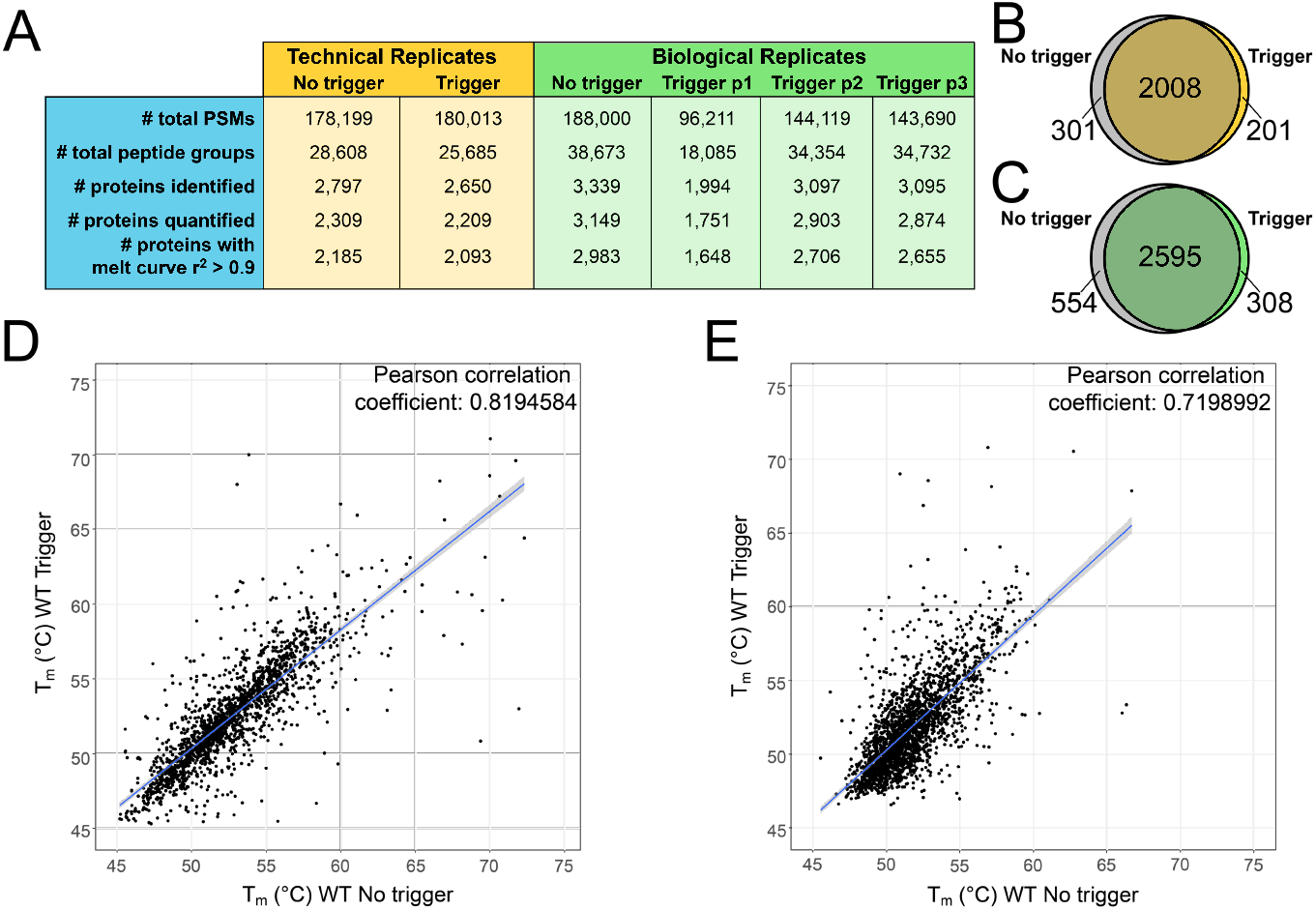
Dataset comparisons from isobaric trigger channel addition. A) Summary of LC-MS/MS data in technical and biological replicates with and without isobaric trigger channel addition. Venn diagrams comparing quantified proteins in no trigger (gray) vs. trigger (yellow/green) in B) technical replicates and C) biological replicate using trigger p2. Correlation plot of the calculated 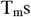 in no trigger vs. trigger in D) technical replicates and E) biological replicates. The blue line represents the linear fit of the data.

A critical feature of mTPP analysis is the ability to accurately calculate melt temperature (T_m_) from the resulting melt curves. To ensure that incorporation of the trigger did not have major impacts on T_m_ calculation of proteins outside of the CPF complex, we performed Pearson correlation analysis of the 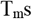 of proteins detected in both the no trigger and trigger samples (Figure 3D, T_m_ data from the TPP package in Supp. Tab. 2). From these we can see a high degree of correlation of 0.82 between the no trigger and trigger samples for proteins which met the criteria for quantitation in our mTPP data analysis workflow (including the number of proteins with melt curves having an r^2^ greater than or equal to 0.9). Additionally, even across biological replicates, there is a strong positive correlation of 0.72 between T_m_ calculations in the no trigger vs. trigger samples (Figure 3E, T_m_ data from the TPP package in Supp. Tab. 2). The ability to make comparisons using biological replicate data would be beneficial in settings with limiting samples where technical replicates may not be feasible in addition to their importance for rigorous statistical analysis.

### An isobaric trigger channel facilitates mTPP analysis of the Cleavage and Polyadenylation Factor Complex

CPF and its accessory factors cleavage factor IA and IB play major roles in RNA processing. CPF is responsible for efficient and specific cleavage and polyadenylation of messenger RNAs ^49, 50^ and has been shown to have important roles in termination of RNA Polymerase II transcription^51, 52^. The CPF complex is currently described as having 14 subunits (Figure 4A) which provide the complex with numerous activities including endonuclease, polyadenylation, and phosphatase functions^53^. Ssu72, which is mutated in the *ssu72-2* yeast strain, is an integral subunit of CPF (Fig. 4A, indicated with a star). Performing mTPP according to the established protocol^28^ resulted in limited detection of CPF (Figure 4C-F, no trigger samples shown in dark/light gray). One notable exception to the low detection of CPF was the subunit Glc7. Along with its presence in CPF, Glc7 is also the catalytic subunit of PP154 and thereby functions in many other protein complexes in eukaryotic cells (reviewed in^55, 56^) where it plays roles in cell cycle regulation and nutrient regulation^54, 57, 58^. Due to these many roles, Glc7 has a higher global abundance than other CPF subunits and is thereby more readily detected.

**Figure 4.**
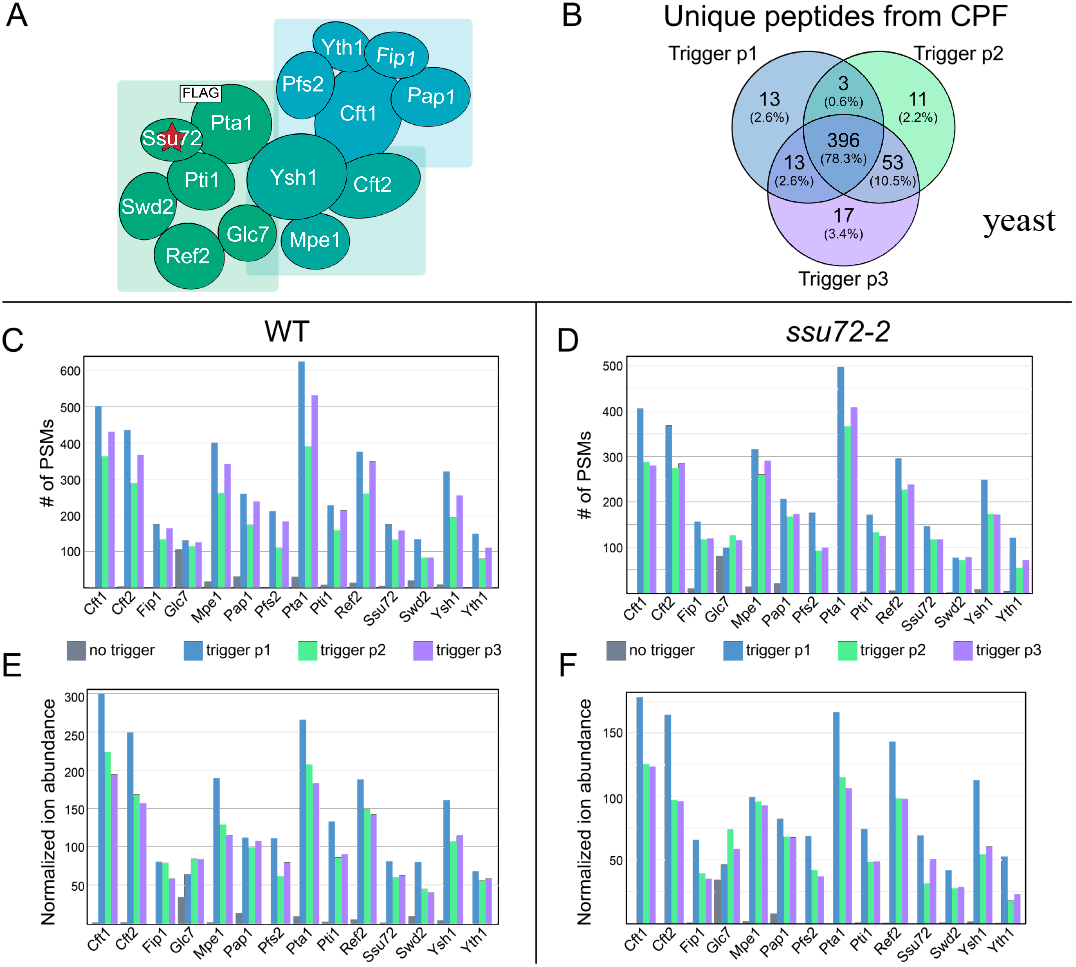
Peptide detection and quantitation for subunits of the Cleavage and Polyadenylation Factor Complex present in the Pta1-FLAG isobaric trigger channel. A) Model of CPF adapted from Casañal *et al* 2017. The red star denotes the mutant protein used in these studies, *ssu72-2;* the white square denotes the FLAG-tagged subunit used for the trigger channel affinity purification, Pta1. B) Venn diagram showing the unique peptides detected for CPF subunits across each WT biological replicate. Number of PSMs for CPF subunits in each C) WT and D) *ssu72-2* replicate experiment. Ion abundanace for CPF subunits normalized to abundance of Pgk1 (x1000) in each E) WT and F) *ssu72-2* replicate experiment.

Previously performed experiments found that the entire CPF complex copurifies with FLAG-tagged Pta1^35^. In theory, addition of an affinity purified CPF sample to one channel of the TMT multiplex would increase the MS1 ion intensity of CPF subunits and would “trigger” the mass spectrometer to pick peptides from CPF complex subunits more often in a DDA analysis than in samples that lack an isobaric trigger. We have previously shown that PSM level detection of affinity purified protein complexes results in highly reproducible quantitation of protein complexes in label-free quantitation workflows ^38, 39^. This prior work found that RNA Polymerase II complex digestions result in the generation of a number of highly detectable peptides and it is likely that this would also be the case for CPF affinity purifications ^39^. If these findings hold true, there should be a significant overlap in unique peptide identifications across the independent LC-MS/MS runs for biological replicates. As shown in Fig. 4B, a significant overlap of unique peptides from CPF complex subunits were identified across the three biological replicates containing the isobaric CPF trigger (peptide data provided in Supp. Tab. 4). Due to the lower overall protein levels in the Trigger p1 sample, a higher level of unique peptide overlap was also observed between Trigger p2 and p3 than was observed between p1/p3 or p1/p2 (Fig. 4B). From an individual subunit perspective, incorporation of the isobaric Pta1-FLAG trigger channel significantly increased identification of most CPF subunits substantially (Figure 3C-F, colored samples). While similar levels of Glc7 were detected across all samples, detection of other complex members was improved significantly in the presence of the isobaric CPF trigger channel. In fact, some CPF subunits that were previously not detected in no trigger samples (such as Cft1, Cft2, and Pfs2) were detected by hundreds of PSMs by utilizing the isobaric CPF trigger channel (Fig. 4C & D). The increased level of PSM detection was accompanied by increased normalized ion abundance (Fig. 4E & F). Overall, this data supports that we can specifically increase reproducible detection and quantitation of proteins of interest for thermal profiling experiments using an isobaric affinity purified trigger channel.

### Mutations in *ssu72-2* do not impact the thermal stability of the CPF protein complex

The CPF complex contains two protein phosphatases, Glc7 and Ssu72. Ssu72 is an integral component of CPF and its function is required for proper termination and 3’-end processing of RNAs ^59–63^. Additionally, its interactions with TFIIB have shown to be critical for the formation of gene loops, which regulate gene expression by linking transcription termination and initiation factors ^64–67^. Much of the characterization of Ssu72 has been accomplished through studies using the *ssu72-2* mutant strain ^41, 59, 65, 68^. The *ssu72-2* TS mutant contains a single mutation, R129A, that confers temperature sensitivity at 37°C. This mutation impairs the catalytic activity of Ssu72, leading to a decrease in transcription elongation efficiency ^41, 68^ and defects in gene looping ^65, 67^. Whether the disrupted phosphatase function in the *ssu72-2* mutant affects the thermal stability of Ssu72 or the CPF complex had not been previously examined.

Detection of CPF with and without the trigger channel resulted in similar numbers of CPF subunits PSMs in *ssu72-2* as in WT which facilitates mTPP analysis of CPF complex thermal stability from a quantitative perspective (Fig. 4C&D). Protein melt curve analysis using the TPP R package (Fig. 5A, mTPP result data in Supp.Tab. 3) showed no obvious changes in any of the 14 CPF subunits in *ssu72-2* relative to WT. Using all biological replicate data, we can define statistically significant changes in protein thermal stability as any ΔT_m_ which falls at least two standard deviations above or below the average ΔT_m_ across the three *ssu72-2* replicates relative to WT. Whole proteome analysis of ΔT_m_ using mTPP found statistically significant decreases in the thermal stability of 59 proteins and increases in the thermal stability of 69 proteins in *ssu72-2* cells (Fig. 5B, Supp. Tab. 5). GO term analysis ^69^ of proteins that had a significant change in thermal stability in *ssu72-2* showed a 2.40-fold enrichment in proteins involved in nucleobase-containing compound biosynthetic process with a p-value of 4.14e^−5^. These results suggest that the defects in transcription caused by disrupted catalytic activity of Ssu72 in this mutant strain are not due to impacts on the stability of Ssu72 or CPF. However, secondary effects of *ssu72-2* functional disruption have been associated with changes in the Nrd1-Nab3-Sen1 complex activity which impact a variety of processes including GTP production ^63, 70, 71^. The temperature sensitivity of this strain is instead likely to be a result of a need for efficient transcription at higher temperatures in order to respond to heat stress^72, 73^. A deeper investigation into the proteins with changes in thermal stability will help to further elucidate the impacts of this catalytic mutant on gene expression.

**Figure 5.**
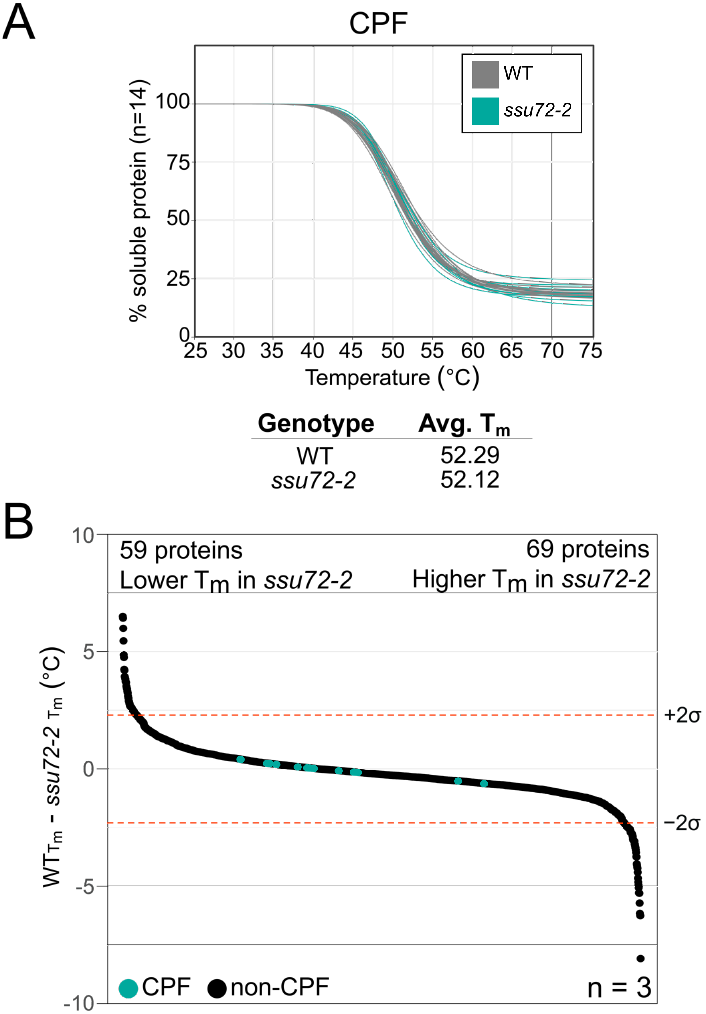
Effects of *ssu72-2* on CPF complex stability and the global proteome. A) mTPP normalized CPF subunit melt curves. Plots for each of the CPF subunits normalized by the TPP package for a representative replicate, Trigger p2. Curves shown in gray are WT and turquoise are *ssu72-2*. Each line represents one of the 14 CPF subunits. Replicates for A are provided in Supp. Fig. 4. B) Waterfall plots visualizing whole proteome changes in melt temperature (T_m_), WT- *ssu72-2*. A total of 2,180 proteins were ordered according to change in T_m_ and plotted. Shown are median values for proteins that were quantified in at least two replicates. Dotted lines signify a confidence interval of 95%. There were significant decreases in thermal stability of 59 proteins and significant increases in thermal stability of 69 proteins. Change in T_m_ and median values provided in Supp. Tab. 5.

## CONCLUSIONS

The integration of an isobaric affinity-purified protein complex trigger channel increased our ability to analyze the low abundance protein complex CPF via mTPP. Our analysis did not observe major effects on the T_m_ estimates of unrelated proteins present in the cell. Protocols for affinity purification would need to be optimized for purity and specificity for optimal use as an isobaric trigger channel. However, since protein complex digestion results in detection of a highly reproducible peptide population, a reasonable alternative approach could include use of a population of purified synthetic peptides or digested recombinant proteins. The use of natively expressed purifications from the system of interest, however, has distinct advantages such as: native protein processing, post-translational modifications, and protein interaction partners.

Use of isobaric purified protein complex trigger channels in TPP studies, and potentially other global proteomics applications, will improve the ability to perform proteomic analysis of low abundance protein complexes and measure systems-level perturbations due to genetic variation(s). The potential for this method to be used across different organisms, even those that are difficult to get large amounts of protein from, is further supported by the adaptation of BASIL for single-cell phosphoproteomics^21^. As many biologically relevant, as well as disease relevant, protein complexes are of relatively low abundance in the cell^74^, improvements in the reproducible detection of such proteins in proteomics experiments would be beneficial to increasing our understanding of the critical cellular mechanisms in normal and disease states.

## Supplementary Material

The supplementary material is available as a PDF and associated XLS tables.

## AUTHOR INFORMATION

### Author Contributions

S.A.P.J.: designed and performed mTPP experiments on biological replicates, analyzed data, prepared the figures, and wrote the manuscript. N.A.M. performed technical replicate mTPP experiments and contributed to the manuscript. J.F.V. affinity purified CPF and confirmed purification via AP-MS (data shown elsewhere). ABW: contributed to the design of experiments. A.L.M.: Oversaw various aspects of the project and provided funding for the project, provided direction on data analysis and figure preparation, and wrote the manuscript. The manuscript was written through contributions of all authors. All authors have given approval to the final version of the manuscript.

### Notes

The authors declare no competing financial interests.

## ACKNOWLEDGMENTS

We would like to thank the current members of the Mosley lab: Whitney Smith-Kinnaman, Katlyn Hughes Burriss, Lynn Bedard, Dominique Baldwin, H.R. Sagara Wijeratne, Gitanjali Roy, and the IUSM proteomics core: Emma Doud and Guihong Qi.

A portion of the funding for this project was provided by National Institute of Health T32 HL007910 (SAPJ) and by the Showalter Research Trust (ALM). NAM was supported in part by the Indiana University Diabetes and Obesity Research Training Program, DeVault Fellowship. This project was supported, in part, with support from the Indiana Clinical and Translational Sciences Institute which is funded by Award Number UL1TR002529 from the National Institutes of Health, National Center for Advancing Translational Sciences, Clinical and Translational Sciences Award. Acquisition of the IUSM Proteomics core instrumentation used for this project was provided by the Indiana University Precision Health Initiative. Some of the TMT reagents were graciously provided via the Thermo Scientific TMT Research Award (SAPJ). The content is solely the responsibility of the authors and does not necessarily represent the official views of the National Institutes of Health.

## REFERENCES

1. Sahni, N.; Yi, S.; Taipale, M.; Fuxman Bass, J. I.; Coulombe-Huntington, J.; Yang, F.; Peng, J.; Weile, J.; Karras, G. I.; Wang, Y.; Kovacs, I. A.; Kamburov, A.; Krykbaeva, I.; Lam, M. H.; Tucker, G.; Khurana, V.; Sharma, A.; Liu, Y. Y.; Yachie, N.; Zhong, Q.; Shen, Y.; Palagi, A.; San-Miguel, A.; Fan, C.; Balcha, D.; Dricot, A.; Jordan, D. M.; Walsh, J. M.; Shah, A. A.; Yang, X.; Stoyanova, A. K.; Leighton, A.; Calderwood, M. A.; Jacob, Y.; Cusick, M. E.; Salehi-Ashtiani, K.; Whitesell, L. J.; Sunyaev, S.; Berger, B.; Barabasi, A. L.; Charloteaux, B.; Hill, D. E.; Hao, T.; Roth, F. P.; Xia, Y.; Walhout, A. J. M.; Lindquist, S.; Vidal, M., Widespread macromolecular interaction perturbations in human genetic disorders. Cell 2015, 161 (3), 647–660.

2. Huttlin, E. L.; Bruckner, R. J.; Paulo, J. A.; Cannon, J. R.; Ting, L.; Baltier, K.; Colby, G.; Gebreab, F.; Gygi, M. P.; Parzen, H.; Szpyt, J.; Tam, S.; Zarraga, G.; Pontano-Vaites, L.; Swarup, S.; White, A. E.; Schweppe, D. K.; Rad, R.; Erickson, B. K.; Obar, R. A.; Guruharsha, K. G.; Li, K.; Artavanis-Tsakonas, S.; Gygi, S. P.; Harper, J. W., Architecture of the human interactome defines protein communities and disease networks. Nature 2017, 545 (7655), 505–509.

3. Chick, J. M.; Munger, S. C.; Simecek, P.; Huttlin, E. L.; Choi, K.; Gatti, D. M.; Raghupathy, N.; Svenson, K. L.; Churchill, G. A.; Gygi, S. P., Defining the consequences of genetic variation on a proteome-wide scale. Nature 2016, 534 (7608), 500–5.

4. Gavin, A. C.; Bosche, M.; Krause, R.; Grandi, P.; Marzioch, M.; Bauer, A.; Schultz, J.; Rick, J. M.; Michon, A. M.; Cruciat, C. M.; Remor, M.; Hofert, C.; Schelder, M.; Brajenovic, M.; Ruffner, H.; Merino, A.; Klein, K.; Hudak, M.; Dickson, D.; Rudi, T.; Gnau, V.; Bauch, A.; Bastuck, S.; Huhse, B.; Leutwein, C.; Heurtier, M. A.; Copley, R. R.; Edelmann, A.; Querfurth, E.; Rybin, V.; Drewes, G.; Raida, M.; Bouwmeester, T.; Bork, P.; Seraphin, B.; Kuster, B.; Neubauer, G.; Superti-Furga, G., Functional organization of the yeast proteome by systematic analysis of protein complexes. Nature 2002, 415 (6868), 141–7.

5. Lambert, J. P.; Ivosev, G.; Couzens, A. L.; Larsen, B.; Taipale, M.; Lin, Z. Y.; Zhong, Q.; Lindquist, S.; Vidal, M.; Aebersold, R.; Pawson, T.; Bonner, R.; Tate, S.; Gingras, A. C., Mapping differential interactomes by affinity purification coupled with data-independent mass spectrometry acquisition. Nat Methods 2013, 10 (12), 1239–45.

6. Go, C. D.; Knight, J. D. R.; Rajasekharan, A.; Rathod, B.; Hesketh, G. G.; Abe, K. T.; Youn, J.-Y.; Samavarchi-Tehrani, P.; Zhang, H.; Zhu, L. Y.; Popiel, E.; Lambert, J.-P.; Coyaud, É.; Cheung, S. W. T.; Rajendran, D.; Wong, C. J.; Antonicka, H.; Pelletier, L.; Raught, B.; Palazzo, A. F.; Shoubridge, E. A.; Gingras, A.-C., A proximity biotinylation map of a human cell. bioRxiv 2019.

7. Rolland, T.; Tasan, M.; Charloteaux, B.; Pevzner, S. J.; Zhong, Q.; Sahni, N.; Yi, S.; Lemmens, I.; Fontanillo, C.; Mosca, R.; Kamburov, A.; Ghiassian, S. D.; Yang, X.; Ghamsari, L.; Balcha, D.; Begg, B. E.; Braun, P.; Brehme, M.; Broly, M. P.; Carvunis, A. R.; Convery-Zupan, D.; Corominas, R.; Coulombe-Huntington, J.; Dann, E.; Dreze, M.; Dricot, A.; Fan, C.; Franzosa, E.; Gebreab, F.; Gutierrez, B. J.; Hardy, M. F.; Jin, M.; Kang, S.; Kiros, R.; Lin, G. N.; Luck, K.; MacWilliams, A.; Menche, J.; Murray, R. R.; Palagi, A.; Poulin, M. M.; Rambout, X.; Rasla, J.; Reichert, P.; Romero, V.; Ruyssinck, E.; Sahalie, J. M.; Scholz, A.; Shah, A. A.; Sharma, A.; Shen, Y.; Spirohn, K.; Tam, S.; Tejeda, A. O.; Trigg, S. A.; Twizere, J. C.; Vega, K.; Walsh, J.; Cusick, M. E.; Xia, Y.; Barabasi, A. L.; Iakoucheva, L. M.; Aloy, P.; De Las Rivas, J.; Tavernier, J.; Calderwood, M. A.; Hill, D. E.; Hao, T.; Roth, F. P.; Vidal, M., A proteome-scale map of the human interactome network. Cell 2014, 159 (5), 1212–1226.

8. Aebersold, R.; Mann, M., Mass-spectrometric exploration of proteome structure and function. Nature 2016, 537 (7620), 347–55.

9. Altelaar, A. F.; Munoz, J.; Heck, A. J., Next-generation proteomics: towards an integrative view of proteome dynamics. Nat Rev Genet 2013, 14 (1), 35–48.

10. Meier, F.; Geyer, P. E.; Virreira Winter, S.; Cox, J.; Mann, M., BoxCar acquisition method enables single-shot proteomics at a depth of 10,000 proteins in 100 minutes. Nature Methods 2018, 15 (6), 440–448.

11. Potel, C. M.; Lin, M.-H.; Heck, A. J. R.; Lemeer, S., Defeating Major Contaminants in Fe3+- Immobilized Metal Ion Affinity Chromatography (IMAC) Phosphopeptide Enrichment. Molecular & Cellular Proteomics 2018, 17 (5), 1028–1034.

12. Humphrey, S. J.; Azimifar, S. B.; Mann, M., High-throughput phosphoproteomics reveals in vivo insulin signaling dynamics. Nature Biotechnology 2015, 33 (9), 990–995.

13. Specht, H.; Slavov, N., Optimizing Accuracy and Depth of Protein Quantification in Experiments Using Isobaric Carriers. J Proteome Res 2020.

14. Slavov, N., Single-cell protein analysis by mass spectrometry. Curr Opin Chem Biol 2020, 60, 1–9.

15. Zhu, Y.; Scheibinger, M.; Ellwanger, D. C.; Krey, J. F.; Choi, D.; Kelly, R. T.; Heller, S.; Barr-Gillespie, P. G., Single-cell proteomics reveals changes in expression during hair-cell development. Elife 2019, 8.

16. Budnik, B.; Levy, E.; Harmange, G.; Slavov, N., SCoPE-MS: mass spectrometry of single mammalian cells quantifies proteome heterogeneity during cell differentiation. Genome Biol 2018, 19 (1), 161.

17. Yi, L.; Tsai, C. F.; Dirice, E.; Swensen, A. C.; Chen, J.; Shi, T.; Gritsenko, M. A.; Chu, R. K.; Piehowski, P. D.; Smith, R. D.; Rodland, K. D.; Atkinson, M. A.; Mathews, C. E.; Kulkarni, R. N.; Liu, T.; Qian, W. J., Boosting to Amplify Signal with Isobaric Labeling (BASIL) Strategy for Comprehensive Quantitative Phosphoproteomic Characterization of Small Populations of Cells. Anal Chem 2019, 91 (9), 5794–5801.

18. McAlister, G. C.; Huttlin, E. L.; Haas, W.; Ting, L.; Jedrychowski, M. P.; Rogers, J. C.; Kuhn, K.; Pike, I.; Grothe, R. A.; Blethrow, J. D.; Gygi, S. P., Increasing the multiplexing capacity of TMTs using reporter ion isotopologues with isobaric masses. Anal Chem 2012, 84 (17), 7469–78.

19. Thompson, A.; Schafer, J.; Kuhn, K.; Kienle, S.; Schwarz, J.; Schmidt, G.; Neumann, T.; Johnstone, R.; Mohammed, A. K.; Hamon, C., Tandem mass tags: a novel quantification strategy for comparative analysis of complex protein mixtures by MS/MS. Anal Chem 2003, 75 (8), 1895–904.

20. Thompson, A.; Wolmer, N.; Koncarevic, S.; Selzer, S.; Bohm, G.; Legner, H.; Schmid, P.; Kienle, S.; Penning, P.; Hohle, C.; Berfelde, A.; Martinez-Pinna, R.; Farztdinov, V.; Jung, S.; Kuhn, K.; Pike, I., TMTpro: Design, Synthesis, and Initial Evaluation of a Proline-Based Isobaric 16-Plex Tandem Mass Tag Reagent Set. Anal Chem 2019, 91 (24), 15941–15950.

21. Tsai, C. F.; Zhao, R.; Williams, S. M.; Moore, R. J.; Schultz, K.; Chrisler, W. B.; Pasa-Tolic, L.; Rodland, K. D.; Smith, R. D.; Shi, T.; Zhu, Y.; Liu, T., An Improved Boosting to Amplify Signal with Isobaric Labeling (iBASIL) Strategy for Precise Quantitative Single-cell Proteomics. Mol Cell Proteomics 2020, 19 (5), 828–838.

22. Chua, X. Y.; Mensah, T.; Aballo, T. J.; Mackintosh, S. G.; Edmondson, R. D.; Salomon, A. R., Tandem Mass Tag approach utilizing pervanadate BOOST channels delivers deeper quantitative characterization of the tyrosine phosphoproteome. Mol Cell Proteomics 2020, mcp.TIR119.0018.

23. Klann, K.; Tascher, G.; Munch, C., Functional Translatome Proteomics Reveal Converging and Dose-Dependent Regulation by mTORC1 and eIF2alpha. Mol Cell 2020, 77 (4), 913–925 e4.

24. Yamamoto, W. R.; Bone, R. N.; Sohn, P.; Syed, F.; Reissaus, C. A.; Mosley, A. L.; Wijeratne, A. B.; True, J. D.; Tong, X.; Kono, T.; Evans-Molina, C., Endoplasmic reticulum stress alters ryanodine receptor function in the murine pancreatic beta cell. J Biol Chem 2019, 294 (1), 168–181.

25. Savitski, M. M.; Reinhard, F. B.; Franken, H.; Werner, T.; Savitski, M. F.; Eberhard, D.; Martinez Molina, D.; Jafari, R.; Dovega, R. B.; Klaeger, S.; Kuster, B.; Nordlund, P.; Bantscheff, M.; Drewes, G., Tracking cancer drugs in living cells by thermal profiling of the proteome. Science 2014, 346 (6205), 1255784.

26. Franken, H.; Mathieson, T.; Childs, D.; Sweetman, G. M.; Werner, T.; Togel, I.; Doce, C.; Gade, S.; Bantscheff, M.; Drewes, G.; Reinhard, F. B.; Huber, W.; Savitski, M. M., Thermal proteome profiling for unbiased identification of direct and indirect drug targets using multiplexed quantitative mass spectrometry. Nat Protoc 2015, 10 (10), 1567–93.

27. Mateus, A.; Kurzawa, N.; Becher, I.; Sridharan, S.; Helm, D.; Stein, F.; Typas, A.; Savitski, M. M., Thermal proteome profiling for interrogating protein interactions. Mol Syst Biol 2020, 16 (3), e9232.

28. Peck Justice, S. A.; Barron, M. P.; Qi, G. D.; Wijeratne, H. R. S.; Victorino, J. F.; Simpson, E. R.; Vilseck, J. Z.; Wijeratne, A. B.; Mosley, A. L., Mutant thermal proteome profiling for characterization of missense protein variants and their associated phenotypes within the proteome. J Biol Chem 2020.

29. Batth, T. S.; Francavilla, C.; Olsen, J. V., Off-line high-pH reversed-phase fractionation for in-depth phosphoproteomics. J Proteome Res 2014, 13 (12), 6176–86.

30. Wang, Y.; Yang, F.; Gritsenko, M. A.; Wang, Y.; Clauss, T.; Liu, T.; Shen, Y.; Monroe, M. E.; Lopez-Ferrer, D.; Reno, T.; Moore, R. J.; Klemke, R. L.; Camp, D. G., 2nd; Smith, R. D., Reversed-phase chromatography with multiple fraction concatenation strategy for proteome profiling of human MCF10A cells. Proteomics 2011, 11 (10), 2019–26.

31. Mertins, P.; Tang, L. C.; Krug, K.; Clark, D. J.; Gritsenko, M. A.; Chen, L.; Clauser, K. R.; Clauss, T. R.; Shah, P.; Gillette, M. A.; Petyuk, V. A.; Thomas, S. N.; Mani, D. R.; Mundt, F.; Moore, R. J.; Hu, Y.; Zhao, R.; Schnaubelt, M.; Keshishian, H.; Monroe, M. E.; Zhang, Z.; Udeshi, N. D.; Mani, D.; Davies, S. R.; Townsend, R. R.; Chan, D. W.; Smith, R. D.; Zhang, H.; Liu, T.; Carr, S. A., Reproducible workflow for multiplexed deep-scale proteome and phosphoproteome analysis of tumor tissues by liquid chromatography-mass spectrometry. Nat Protoc 2018, 13 (7), 1632–1661.

32. Hogrebe, A.; von Stechow, L.; Bekker-Jensen, D. B.; Weinert, B. T.; Kelstrup, C. D.; Olsen, J. V., Benchmarking common quantification strategies for large-scale phosphoproteomics. Nat Commun 2018, 9 (1), 1045.

33. Gilar, M.; Olivova, P.; Daly, A. E.; Gebler, J. C., Orthogonality of separation in two-dimensional liquid chromatography. Anal Chem 2005, 77 (19), 6426–34.

34. Ludwig, K. R.; Schroll, M. M.; Hummon, A. B., Comparison of In-Solution, FASP, and S-Trap Based Digestion Methods for Bottom-Up Proteomic Studies. J Proteome Res 2018, 17 (7), 2480–2490.

35. Victorino, J. F.; Fox, M. J.; Smith-Kinnaman, W. R.; Peck Justice, S. A.; Burriss, K. H.; Boyd, A. K.; Zimmerly, M. A.; Chan, R. R.; Hunter, G. O.; Liu, Y.; Mosley, A. L., RNA Polymerase II CTD phosphatase Rtr1 fine-tunes transcription termination. PLoS Genet 2020, 16 (3), e1008317.

36. Bedard, L. G.; Dronamraju, R.; Kerschner, J. L.; Hunter, G. O.; Axley, E. D.; Boyd, A. K.; Strahl, B. D.; Mosley, A. L., Quantitative Analysis of Dynamic Protein Interactions during Transcription Reveals a Role for Casein Kinase II in Polymerase-associated Factor (PAF) Complex Phosphorylation and Regulation of Histone H2B Monoubiquitylation. J Biol Chem 2016, 291 (26), 13410–20.

37. Smith-Kinnaman, W. R.; Berna, M. J.; Hunter, G. O.; True, J. D.; Hsu, P.; Cabello, G. I.; Fox, M. J.; Varani, G.; Mosley, A. L., The interactome of the atypical phosphatase Rtr1 in Saccharomyces cerevisiae. Mol Biosyst 2014, 10 (7), 1730–41.

38. Mosley, A. L.; Hunter, G. O.; Sardiu, M. E.; Smolle, M.; Workman, J. L.; Florens, L.; Washburn, M. P., Quantitative proteomics demonstrates that the RNA polymerase II subunits Rpb4 and Rpb7 dissociate during transcriptional elongation. Mol Cell Proteomics 2013, 12 (6), 1530–8.

39. Mosley, A. L.; Sardiu, M. E.; Pattenden, S. G.; Workman, J. L.; Florens, L.; Washburn, M. P., Highly reproducible label free quantitative proteomic analysis of RNA polymerase complexes. Mol Cell Proteomics 2011, 10 (2), M110 000687.

40. McGinty, R. J.; Puleo, F.; Aksenova, A. Y.; Hisey, J. A.; Shishkin, A. A.; Pearson, E. L.; Wang, E. T.; Housman, D. E.; Moore, C.; Mirkin, S. M., A Defective mRNA Cleavage and Polyadenylation Complex Facilitates Expansions of Transcribed (GAA)n Repeats Associated with Friedreich’s Ataxia. Cell Rep 2017, 20 (10), 2490–2500.

41. Pappas, D. L.; Hampsey, M., Functional Interaction between Ssu72 and the Rpb2 Subunit of RNA Polymerase II in Saccharomyces cerevisiae. 2000, 20 (22), 8343–8351.

42. Funakoshi, M.; Hochstrasser, M., Small epitope-linker modules for PCR-based C-terminal tagging inSaccharomyces cerevisiae. Yeast 2009, 26 (3), 185–192.

43. Perez-Riverol, Y.; Csordas, A.; Bai, J.; Bernal-Llinares, M.; Hewapathirana, S.; Kundu, D. J.; Inuganti, A.; Griss, J.; Mayer, G.; Eisenacher, M.; Pérez, E.; Uszkoreit, J.; Pfeuffer, J.; Sachsenberg, T.; Yılmaz, Ş.; Tiwary, S.; Cox, J.; Audain, E.; Walzer, M.; Jarnuczak, A. F.; Ternent, T.; Brazma, A.; Vizcaíno, J. A., The PRIDE database and related tools and resources in 2019: improving support for quantification data. Nucleic Acids Research 2019, 47 (D1), D442–D450.

44. Oliveros, J. C., Venny. An interactive tool for comparing lists with Venn’s diagrams. 2007–2015.

45. Wickham, H. ggplot2: Elegant Graphics for Data Analysis, Springer-Verlag New York: 2016.

46. Childs D, K. N., Franken H, Doce C, Savitski M, Huber W TPP: Analyze thermal proteome profiling (TPP) experiments, 3.10.0; 2018.

47. Walmsley, S. J.; Rudnick, P. A.; Liang, Y.; Dong, Q.; Stein, S. E.; Nesvizhskii, A. I., Comprehensive analysis of protein digestion using six trypsins reveals the origin of trypsin as a significant source of variability in proteomics. J Proteome Res 2013, 12 (12), 5666–80.

48. Burkhart, J. M.; Schumbrutzki, C.; Wortelkamp, S.; Sickmann, A.; Zahedi, R. P., Systematic and quantitative comparison of digest efficiency and specificity reveals the impact of trypsin quality on MS-based proteomics. J Proteomics 2012, 75 (4), 1454–62.

49. Chen, J.; Moore, C., Separation of factors required for cleavage and polyadenylation of yeast pre-mRNA. 1992, 12 (8), 3470–3481.

50. Kessler, M. M.; Zhao, J.; Moore, C. L., Purification of the Saccharomyces cerevisiae cleavage/polyadenylation factor I. Separation into two components that are required for both cleavage and polyadenylation of mRNA 3’ ends. J Biol Chem 1996, 271 (43), 27167–75.

51. Proudfoot, N. J., Transcriptional termination in mammals: Stopping the RNA polymerase II juggernaut. Science 2016, 352 (6291), aad9926.

52. Eaton, J. D.; Davidson, L.; Bauer, D. L. V.; Natsume, T.; Kanemaki, M. T.; West, S., Xrn2 accelerates termination by RNA polymerase II, which is underpinned by CPSF73 activity. Genes Dev 2018, 32 (2), 127–139.

53. Casanal, A.; Kumar, A.; Hill, C. H.; Easter, A. D.; Emsley, P.; Degliesposti, G.; Gordiyenko, Y.; Santhanam, B.; Wolf, J.; Wiederhold, K.; Dornan, G. L.; Skehel, M.; Robinson, C. V.; Passmore, L. A., Architecture of eukaryotic mRNA 3’-end processing machinery. Science 2017, 358 (6366), 1056–1059.

54. Feng, Z. H.; Wilson, S. E.; Peng, Z. Y.; Schlender, K. K.; Reimann, E. M.; Trumbly, R. J., The Yeast Glc7-Gene Required for Glycogen Accumulation Encodes a Type-1 Protein Phosphatase. Journal of Biological Chemistry 1991, 266 (35), 23796–23801.

55. Martín, R.; Stonyte, V.; Lopez-Aviles, S., Protein Phosphatases in G1 Regulation. International Journal of Molecular Sciences 2020, 21 (2), 395.

56. Moura, M.; Conde, C., Phosphatases in Mitosis: Roles and Regulation. Biomolecules 2019, 9 (2), 55.

57. Tu, J.; Carlson, M., The GLC7 type 1 protein phosphatase is required for glucose repression in Saccharomyces cerevisiae. Mol Cell Biol 1994, 14 (10), 6789–96.

58. Ramaswamy, N. T.; Li, L.; Khalil, M.; Cannon, J. F., Regulation of yeast glycogen metabolism and sporulation by Glc7p protein phosphatase. Genetics 1998, 149 (1), 57–72.

59. Dichtl, B.; Blank, D.; Ohnacker, M.; Friedlein, A.; Roeder, D.; Langen, H.; Keller, W., A Role for SSU72 in Balancing RNA Polymerase II Transcription Elongation and Termination. Molecular Cell 2002, 10 (5), 1139–1150.

60. Nedea, E.; He, X.; Kim, M.; Pootoolal, J.; Zhong, G.; Canadien, V.; Hughes, T.; Buratowski, S.; Moore, C. L.; Greenblatt, J., Organization and Function of APT, a Subcomplex of the Yeast Cleavage and Polyadenylation Factor Involved in the Formation of mRNA and Small Nucleolar RNA 3’-Ends. 2003, 278 (35), 33000–33010.

61. He, X.; Khan, A. U.; Cheng, H.; Pappas, D. L., Jr.; Hampsey, M.; Moore, C. L., Functional interactions between the transcription and mRNA 3’ end processing machineries mediated by Ssu72 and Sub1. Genes Dev 2003, 17 (8), 1030–42.

62. Steinmetz, E. J.; Brow, D. A., Ssu72 Protein Mediates Both Poly(A)-Coupled and Poly(A)-Independent Termination of RNA Polymerase II Transcription. 2003, 23 (18), 6339–6349.

63. Zhang, D. W.; Mosley, A. L.; Ramisetty, S. R.; Rodriguez-Molina, J. B.; Washburn, M. P.; Ansari, A. Z., Ssu72 phosphatase-dependent erasure of phospho-Ser7 marks on the RNA polymerase II C-terminal domain is essential for viability and transcription termination. J Biol Chem 2012, 287 (11), 8541–51.

64. Ansari, A.; Hampsey, M., A role for the CPF 3’-end processing machinery in RNAP II-dependent gene looping. Genes Dev 2005, 19 (24), 2969–78.

65. Allepuz-Fuster, P.; O’Brien, M. J.; Gonzalez-Polo, N.; Pereira, B.; Dhoondia, Z.; Ansari, A.; Calvo, O., RNA polymerase II plays an active role in the formation of gene loops through the Rpb4 subunit. Nucleic Acids Res 2019, 47 (17), 8975–8987.

66. Singh, B. N.; Hampsey, M., A transcription-independent role for TFIIB in gene looping. Mol Cell 2007, 27 (5), 806–16.

67. Tan-Wong, S. M.; Zaugg, J. B.; Camblong, J.; Xu, Z.; Zhang, D. W.; Mischo, H. E.; Ansari, A. Z.; Luscombe, N. M.; Steinmetz, L. M.; Proudfoot, N. J., Gene loops enhance transcriptional directionality. Science 2012, 338 (6107), 671–5.

68. Reyes-Reyes, M.; Hampsey, M., Role for the Ssu72 C-terminal domain phosphatase in RNA polymerase II transcription elongation. Mol Cell Biol 2007, 27 (3), 926–36.

69. Mi, H.; Huang, X.; Muruganujan, A.; Tang, H.; Mills, C.; Kang, D.; Thomas, P. D., PANTHER version 11: expanded annotation data from Gene Ontology and Reactome pathways, and data analysis tool enhancements. Nucleic Acids Res 2017, 45 (D1), D183’D189.

70. Ganem, C.; Devaux, F.; Torchet, C.; Jacq, C.; Quevillon-Cheruel, S.; Labesse, G.; Facca, C.; Faye, G., Ssu72 is a phosphatase essential for transcription termination of snoRNAs and specific mRNAs in yeast. EMBO J 2003, 22 (7), 1588–98.

71. Loya, T. J.; O’Rourke, T. W.; Reines, D., A genetic screen for terminator function in yeast identifies a role for a new functional domain in termination factor Nab3. Nucleic Acids Res 2012, 40 (15), 7476–91.

72. Mahat, D. B.; Salamanca, H. H.; Duarte, F. M.; Danko, C. G.; Lis, J. T., Mammalian Heat Shock Response and Mechanisms Underlying Its Genome-wide Transcriptional Regulation. Mol Cell 2016, 62 (1), 63–78.

73. Duarte, F. M.; Fuda, N. J.; Mahat, D. B.; Core, L. J.; Guertin, M. J.; Lis, J. T., Transcription factors GAF and HSF act at distinct regulatory steps to modulate stress-induced gene activation. Genes Dev 2016, 30 (15), 1731–46.

74. Ho, B.; Baryshnikova, A.; Brown, G. W., Unification of Protein Abundance Datasets Yields a Quantitative Saccharomyces cerevisiae Proteome. Cell Syst 2018, 6 (2), 192–205 e3.

