## Supplementary material for "Boosting detection of low abundance proteins in thermal proteome profiling experiments by addition of an isobaric trigger channel to TMT multiplexes": Supp. Fig.

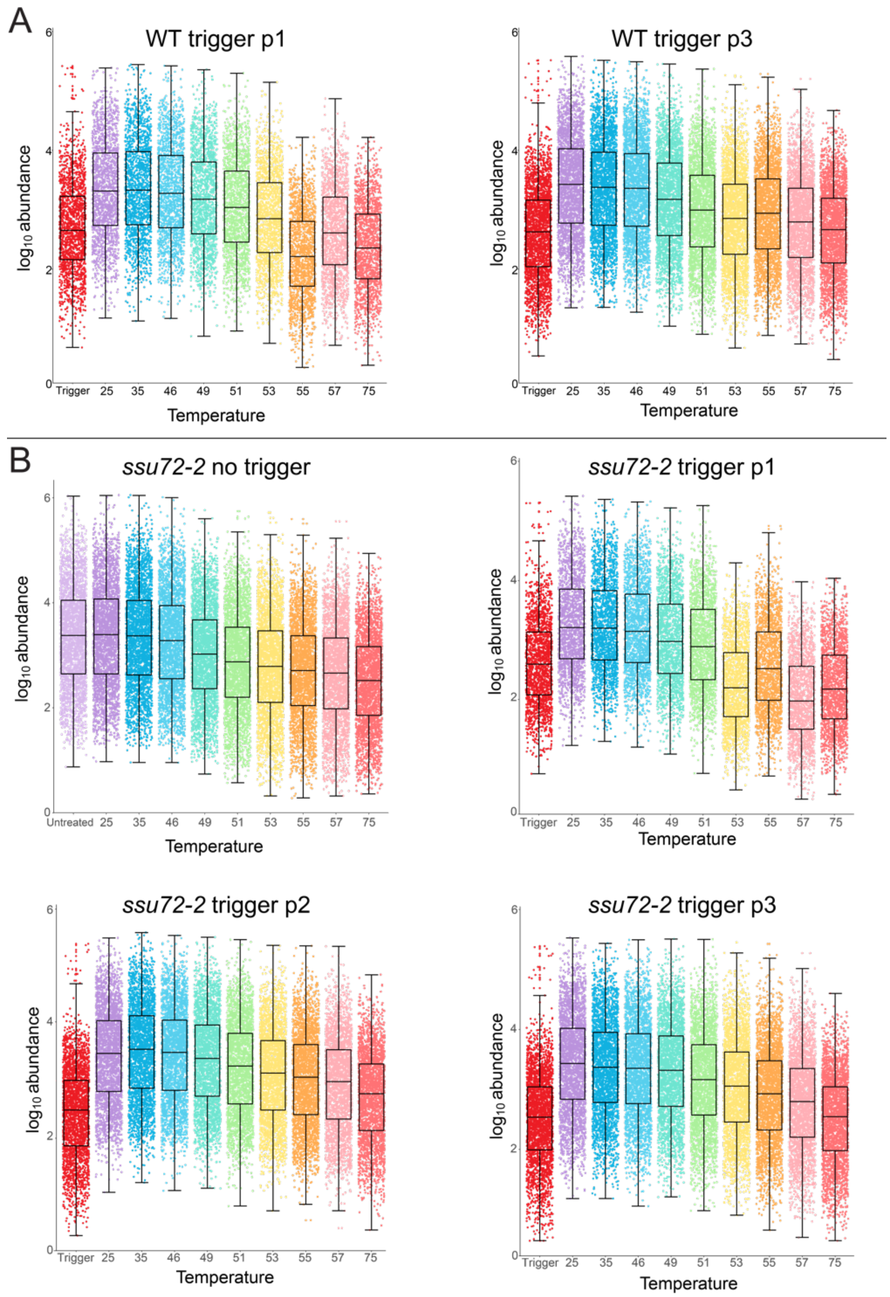


**Supplemental Figure 1: Additional protein abundance dot plots**

Dot plots of protein abundance values for every protein detected in A) WT trigger replicates p1 and p3 and B) *ssu72-2* no trigger and trigger replicates p1, p2, and p3

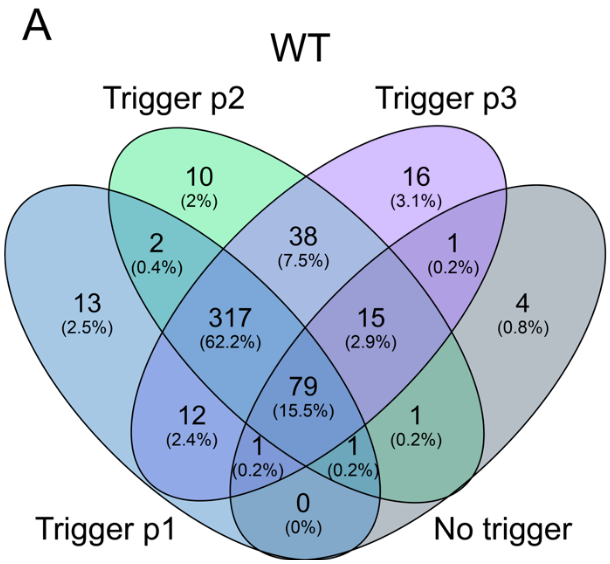


**Supplemental Figure 2: Unique CPF peptide detection**

Venn diagrams of the overlap in peptides detected for each of the 14 CPF subunits across each experiment in A) WT and B) *ssu72-2*.

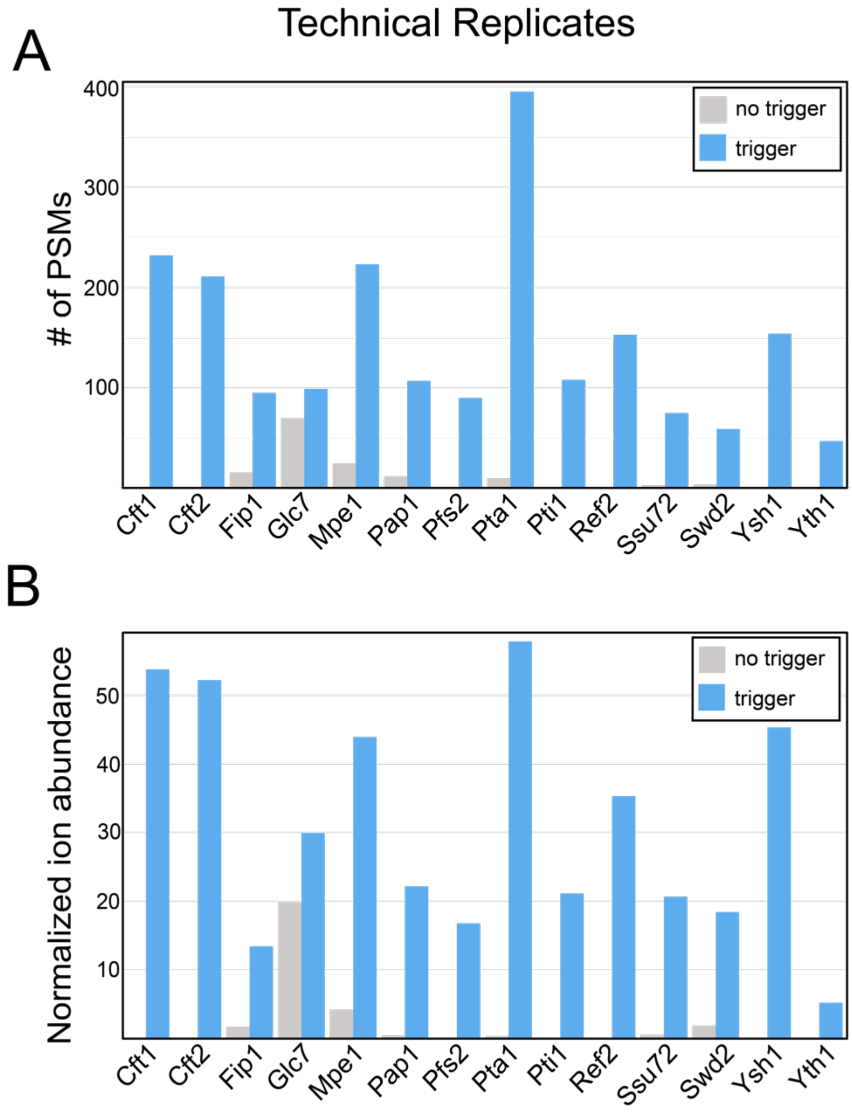


**Supplemental Figure 3: PSMs and ion abundance measurements for CPF subunits in WT technical replicates**
A) Numbers of PSMs and B) ion abundance normalized to the abundance of Pgk1 for each CPF subunit in WT

A

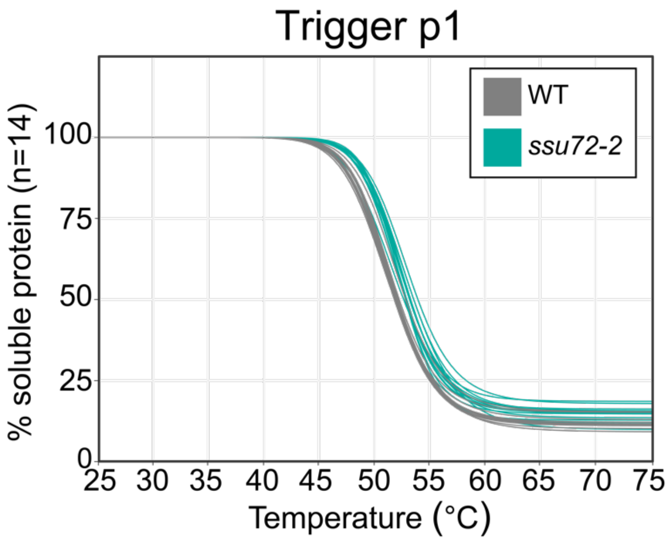


B

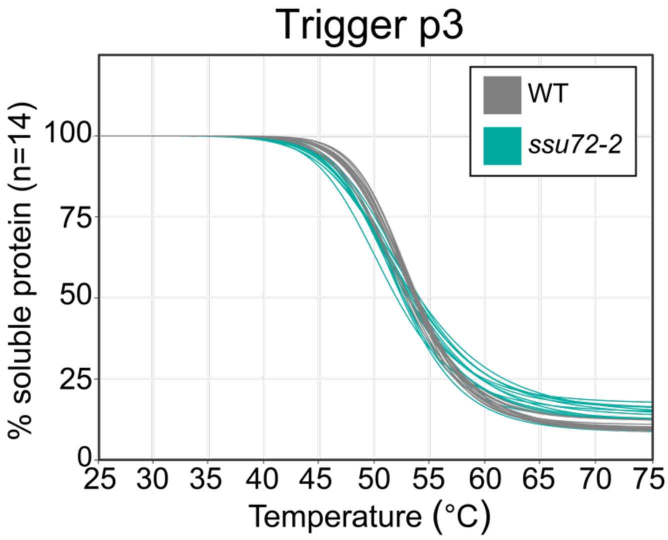


**Supplemental Figure 4: Replicate CPF melt curves**
mTPP normalized CPF subunit melt curves. Plots for each of the CPF subunits normalized by the TPP package for trigger p1 and p3; trigger p2 is shown in Figure 5. Curves shown in gray are WT and turquoise are *ssu72-2*. Each line represents one of the 14 CPF subunits.

**Supporting Table 1: mTPP data**
Table of data obtained from mTPP experiments exported from Proteome Discoverer. Each genotype and replicate are provided as a separate sheet within the document. Provided is the accession number, protein description from Uniprot, Sum PEP score, percent coverage, number of peptides, number of peptide spectral matches (PSMs), number of unique peptides, number of amino acids, molecular weight, raw abundances for each channel, and percent of soluble protein as normalized to the lowest heat-treated sample (35°).

**Supporting Table 2: TPP package results, WT no trigger vs WT trigger**
Output results provided from the TPP package of the comparing WT no trigger vs WT trigger in technical and biological replicates. Input data provided in Table 1.

**Supporting Table 3: TPP package results, WT vs. *ssu72-2***
Output results provided from the TPP package comparing WT vs. *ssu72-2* in one no trigger biological replicate and three trigger biological replicates.

**Supporting Table 4: Peptide groups for CPF subunits in WT and *ssu72-2***
Table of unique peptide groups in WT and *ssu72-2* exported from Proteome Discoverer. Each genotype and replicate are provided as a separate sheet within the document.

**Supporting Table 5: Changes in Tm and median changes**
Table of change in $T_m$ (calculated from the results in Table 3) and median change in $T_m$. $T_m$ values for *ssu72-2* were subtracted from WT to get changes in $T_m$. Median values were calculated for each protein that was quantified and was given a $T_m$ and >0.9 $r^2$ by the TPP package in at least two replicates.
